# Composition dependent phase separation underlies directional flux through the nucleolus

**DOI:** 10.1101/809210

**Authors:** Joshua A. Riback, Lian Zhu, Mylene C. Ferrolino, Michele Tolbert, Diana M. Mitrea, David W. Sanders, Ming-Tzo Wei, Richard W. Kriwacki, Clifford P. Brangwynne

**Affiliations:** Department of Chemical and Biological Engineering, Princeton University, Princeton, NJ 08544, USA; Department of Structural Biology, St. Jude Children’s Research Hospital, Memphis, TN 38103, USA; Lewis Sigler Institute for Integrative Genomics, Princeton University, Princeton, NJ 08544, USA; Howard Hughes Medical Institute

**Author notes:** Equal contribution (Joshua A. Riback, Lian Zhu).

## Abstract

Intracellular bodies such as nucleoli, Cajal bodies, and various signaling assemblies, represent membraneless organelles, or condensates, that form via liquid-liquid phase separation (LLPS)^1,2^. Biomolecular interactions, particularly homotypic interactions mediated by self-associating intrinsically disordered protein regions (IDRs), are thought to underlie the thermodynamic driving forces for LLPS, forming condensates that can facilitate the assembly and processing of biochemically active complexes, such as ribosomal subunits within the nucleolus. Simplified model systems^3–6^ have led to the concept that a single fixed saturation concentration (C_sat_) is a defining feature of endogenous LLPS^7–9^, and has been suggested as a mechanism for intracellular concentration buffering^2,7,8,10^. However, the assumption of a fixed C_sat_ remains largely untested within living cells, where the richly multicomponent nature of condensates could complicate this simple picture. Here we show that heterotypic multicomponent interactions dominate endogenous LLPS, and give rise to nucleoli and other condensates that do not exhibit a fixed C_sat_. As the concentration of individual components is varied, their partition coefficients change, in a manner that can be used to extract thermodynamic interaction energies, that we interpret within a framework we term polyphasic interaction thermodynamic analysis (PITA). We find that heterotypic interactions between protein and RNA components stabilize a variety of archetypal intracellular condensates, including the nucleolus, Cajal bodies, stress granules, and P bodies. These findings imply that the composition of condensates is finely tuned by the thermodynamics of the underlying biomolecular interaction network. In the context of RNA processing condensates such as the nucleolus, this stoichiometric self-tuning manifests in selective exclusion of fully-assembled RNP complexes, providing a thermodynamic basis for vectorial ribosomal RNA (rRNA) flux out of the nucleolus. The PITA methodology is conceptually straightforward and readily implemented, and it can be broadly utilized to extract thermodynamic parameters from microscopy images. These approaches pave the way for a deep understanding of the thermodynamics of multi-component intracellular phase behavior and its interplay with nonequilibrium activity characteristic of endogenous condensates.

---

To determine the thermodynamics of LLPS for intracellular condensates we first focused on the liquid granular component (GC) of nucleoli within HeLa cells, in particular on the protein Nucleophosmin (NPM1), which is known to be a key driver of nucleolar phase separation^11,12^. Under typical endogenous expression levels, we estimate NPM1 concentration in the nucleoplasm to be roughly C^dil^~4μM; from simple binary phase separation models (i.e. Regular solution theory)^13^, this apparent saturation concentration, C_sat_, is expected to be fixed even under varied NPM1 expression levels (**Fig. 1C**, **Supplemental Text**). Consistent with previous studies^11^, overexpression of NPM1 resulted in larger nucleoli, underscoring the importance of NPM1 in nucleolar assembly (**Fig. 1A**). However, with these increased levels of NPM1, the nucleoplasmic concentration did not remain fixed at a single C_sat_, but instead increased by roughly 10-fold (**Fig. 1B and Supplemental Text**). Interestingly, the NPM1 concentration within the dense phase nucleolus, C^den^, also increases, but the ratio of the dense to dilute concentrations, known as the partition coefficient K=C^den^/C^dil^, decreased significantly (**Fig. S1**).

**Figure 1.**
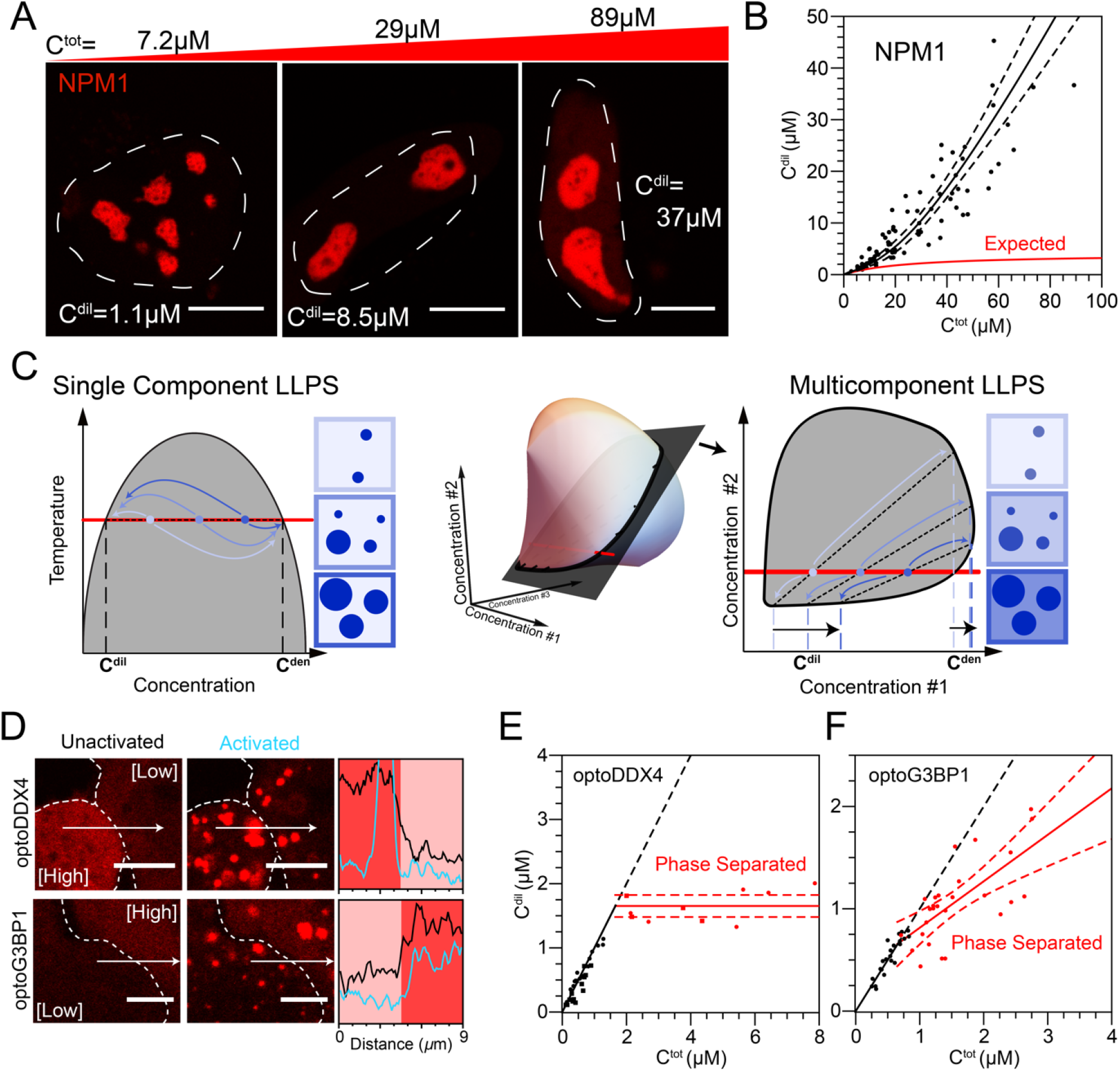
Multicomponent LLPS results in elimination of a fixed C_sat_ and the emergence of a concentration-dependent phase stability. (A) Images of cells expressing NPM1-mCherry denoting the total nuclear concentration (C^tot^) and the nucleoplasmic concentration (C^dil^) of NPM1-mCherry in the top and within the image, respectively. The white dashed lines denote the nuclear boundary as defined by NPM1 signal. Scale bar is 10 microns. (B) Concentration of NPM1-mCherry in the nucleoplasm (C^dil^) with respect to the total NPM1-mCherry concentration in the nucleus (C^tot^). The expected trend for a single C_sat_ is shown in red as described in the text. (C) Graphical representation of phase diagrams for both single and multicomponent LLPS showing fixed and non-fixed C_sat_, respectively. (D) OptoDroplet constructs with optoDDX4 (top row and E) or optoG3BP1 (bottom row and F) with the total and cytoplasmic (circles)/nucleoplasmic (squares) concentrations in a cell before and after light activation, respectively. OptoG3BP1 experiments are arsenite-stressed cells with G3BP1A/B knocked out; optoDDX4 data reproduced from^14^. Here scale bars are 5 microns. Line scans shown correspond to intensity traces before and after activation in black and blue, respectively.

To elucidate the underlying biophysics of this observed non-fixed C_sat_ within living cells, we examined phase separation of model biomimetic condensates not natively present within the cell. We took advantage of the optoDroplet system^4^, developed for controlling intracellular phase separation, by fusing the blue-light-dependent oligomerizing protein Cry2 to the IDR of DDX4, which drives phase separation of exogenous condensates through predominately homotypic interactions^3,4,10^. Consistent with our prior work^14^, at total cellular concentrations above ~1.7μM, light activates droplet formation, and nucleoplasmic and cytoplasmic C^dil^ remains at a fixed value, suggesting a fixed C_sat_~1.7μM (**Fig. 1D-E**). We next asked whether a fixed C_sat_ would be observed with light induction of stress granules, multi-component, stress-inducible condensates which assemble through heterotypic protein-mRNA interactions^15^. We replaced the oligomerization domain of G3BP1, a critical stress granule protein, with Cry2, and expressed this construct in G3BP1/2 knockout cells under arsenite stress. At total cytoplasmic concentrations above ~0.7μM, light triggers droplet formation; however, unlike the synthetic DDX4 case, the C^dil^ was not fixed, but instead increased with increased total concentrations (**Fig. 1D, F**), similar to the behavior of NPM1 (**Fig. 1A,B**). These results are not restricted to light-induced oligomerization of G3BP1 using the optogenetic system, as increasing expression of G3BP1 in a G3BP1/2 KO cell line exhibits a similar increase in the C^dil^ (**Fig. S2**).

These data suggest that multicomponent condensates are not governed by a fixed C_sat_, as expected for a single component binodal phase boundary at fixed temperature, but instead governed by a higher dimensional phase diagram. Indeed, for multicomponent systems, the Gibbs phase rule dictates that the concentrations 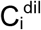 and 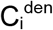 for a component i, will depend on the concentrations of all other components^16^ (**Fig. 1C**). To probe this concentration-dependent thermodynamics, we increase the concentration of a biomolecule *in vivo* or *in vitro*, altering stoichiometry to shift from a heterotypic to a homotypic composition of the system (**Fig. 2A**). This changes the apparent partition coefficient, allowing us to quantify the transfer free energy of component i from the dilute to the dense phase, 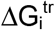 (**Fig. 2B**); a thermodynamic derivation yields the relationship, 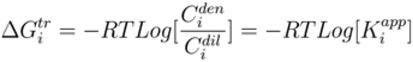 **Supplemental Text**). For components which contribute to phase separation (e.g. act to scaffold the condensate meshwork), their transfer free energy reports on the stability of interactions driving phase separation. Validation of the compositional dependence of LLPS can be understood based on theoretical grounds (**Supplemental Text**) and quantitatively using analytical methods such as Flory-Huggins theory (**Fig. S3**).

**Figure 2.**
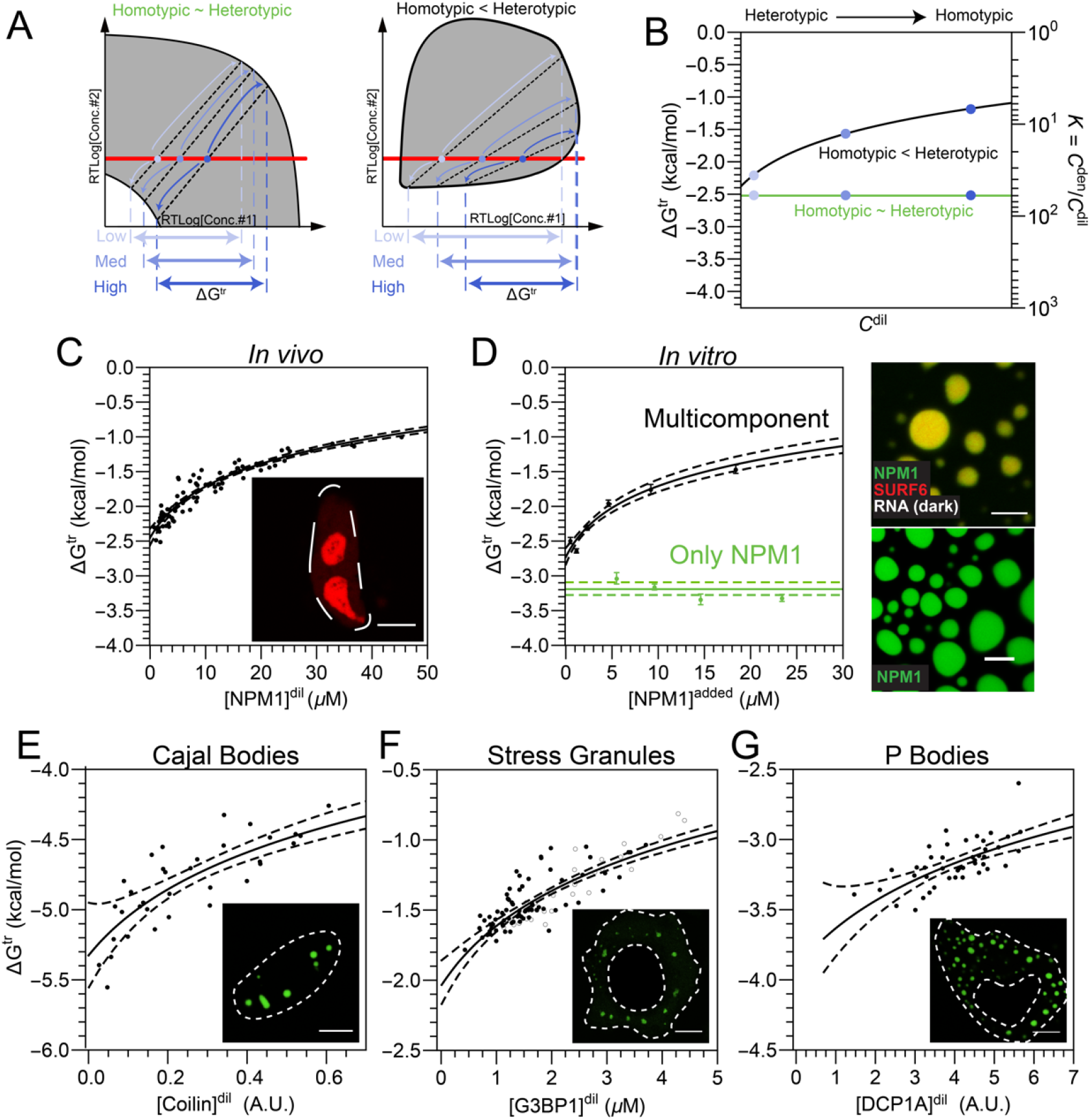
Determining the contribution of heterotypic and homotypic interactions driving condensate formation *in vivo* and *in vitro*. (A) Schematic illustrating the connection between the phase diagram and the transfer free energy of a component when heterotypic interactions are equal (left) to or stronger (right) compared to homotypic interactions. (B) Accompanying schematic detailing the qualitative change in the transfer free energy of component 1 with an increase in its expression for the two cases in (A). (C) Thermodynamic dependence of NPM1-mCherry transfer from the nucleoplasm into the nucleolus GC, as a function of its increased expression (concentration in the nucleoplasm). Inset, image from Fig 1A to highlight that these data represent a reanalysis of those experiments. (D) In vitro reconstitution experiments showing change in the transfer free energy for NPM1 as a function of added NPM1. Image of NPM1 droplets with 5% PEG (bottom right) and of ternary NPM1:SURF6N:rRNA droplets in buffer (top right). Change in transfer free energy for Coilin-EYFP (E), G3BP1 (F, −GFP empty, −mCherry filled), and DCP1A-EYFP (G) from the dilute phase (i.e. nucleoplasm or cytoplasm) to Cajal bodies, arsenite-induced Stress granules, and P-bodies (i.e. dense phases), respectively. All scale bars are 10 microns.

Applying this framework to our NPM1 results (**Fig. 1A-B**), as the NPM1 concentration is increased, the partition coefficient of NPM1 into the nucleolus decreases (**Fig. S1B**), and thus the transfer free energy ΔG^tr^ for NPM1 between the condensed and dilute phases becomes less negative, and thus destabilizing; this destabilizing effect at higher NPM1 concentrations implies that heterotypic, rather than homotypic (*i.e.* NPM1-NPM1), interactions drive nucleolar assembly (**Fig. 2C**). To further test this conclusion, we focused on *in vitro* reconstitution of the nucleolar GC. In addition to NPM1, key GC components include ribosomal RNA (rRNA), and multivalent poly-arginine motif-containing proteins (Arg-proteins) such as SURF6, and ribosomal proteins (r-proteins). Utilizing a well-established system for GC phase separation *in vitro* ^11,12,17,18^, we formed either NPM1-only droplets with 5% PEG as a crowder (**Fig. 2D**, **bottom image**) or multicomponent droplets containing NPM1, the N-terminus of SURF6 (SURF6N), and rRNA (**Fig. 2D, upper image)**. As expected for single-component phase separation, as more NPM1 was added to the NPM1-only droplets, the transfer free energy remained roughly constant (**Fig. 2D**, **green data**). In contrast, for multicomponent droplets, the transfer free energy became significantly less negative (i.e. destabilizing) as more NPM1 was added, exactly as observed in living cells (**Fig. 2D**, **black data**). Remarkably, similar behavior was observed with a number of different intracellular condensates and their associated key scaffolding proteins: Coilin in Cajal bodies, G3BP1 in arsenite-triggered stress granules, and DCP1A in P-Bodies (**Fig. 2E-G**); in each of these cases, the transfer free energy also became less negative with increasing protein concentrations. These data contrast with the view that condensates are stabilized by predominantly homotypic interactions, for example those mediated by self-associating IDRs, and instead suggest that heterotypic interactions provide the dominant internal cohesivity stabilizing LLPS, not only for nucleoli but also for other intracellular condensates.

We next probed which heterotypic interactions drive phase separation of the nucleolus, by monitoring one component’s transfer free energy while changing the concentration of another (**Fig. 3A**). In our multicomponent in vitro mimic, we find that increasing NPM1 or SURF6N destabilizes the transfer of SURF6N (**Fig. S4**), again consistent with heterotypic interactions driving SURF6 to nucleoli. In living cells, SURF6 also exhibits behavior similar to NPM1, with a destabilizing increase in the transfer free energy observed with increasing SURF6 concentration (**Fig. 3B**, black lines). Interestingly, this in vivo destabilization is dramatically amplified with increasing NPM1 concentrations (**Fig. 3B**). From these data, we determined the change in the transfer free energy of SURF6 as a function of NPM1 by referencing to the energy expected without NPM1 overexpression, *i.e* 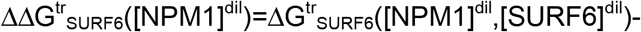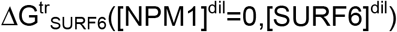. Remarkably, plotting 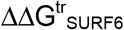. NPM1 collapses the data onto a single master curve (**Fig. 3D**, **Supplemental Text**), highlighting a tight thermodynamic linkage between NPM1 and SURF6. This behavior contrasts with that of ribosomal proteins (r-proteins), which exhibit strong and specific rRNA binding, and exhibit a transfer free energy which is statistically insensitive to NPM1 concentrations (**Fig. 3D and S5**).

**Figure 3.**
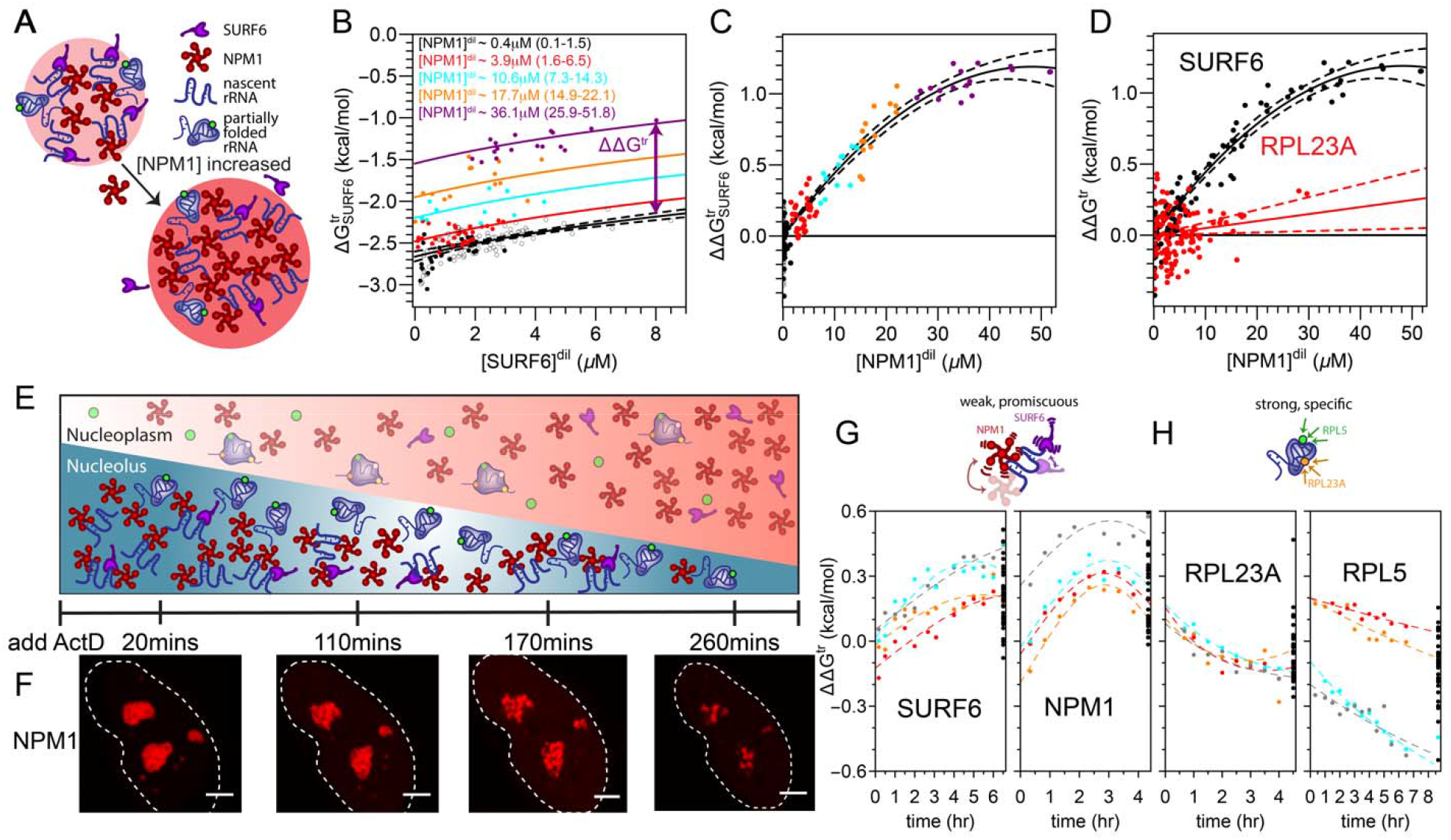
Heterotypic interactions between nucleolar proteins and rRNA underly nucleolar thermodynamics. (A) Schematic of proposed mechanism for the dilution of non-NPM1 molecular interactions in the dense phase due to NPM1 overexpression. Only relevant species shown for clarity. (B) Change in the transfer free energy of *in vivo* SURF6 with overexpression of NPM1 plotted against SURF6 concentration (colors are different concentrations of NPM1 in μM as indicated with mean and range values, open circles are cells without additional NPM1 expressed). The method of calculating ΔΔG^tr^ at a referenced nucleoplasmic SURF6 concentration is shown via arrows and displaced lines in (B) and (C) shown as a function of NPM1 concentration with the same colored concentrations as B. (D) Comparison between the change in ΔΔG^tr^ with NPM1 overexpression for SURF6 and RPL23A as indicated. (E) Schematic of ActD treatment on the GC of nucleoli with time. (F) Images of one of the five cells at indicated times post ActD treatment. Corresponding quantification for NPM1 cells is shown in Fig. S5. Scale bar is 5 microns. Dependence of ΔΔG^tr^ of (G) NPM1 and SURF6 and (H) RPL23A and RPL5 with ActD treatment. Each color for each plot represents an individual cell followed with time. Black points are an ensemble of cells (n>20) measured at the indicated time points. Schematics highlighting the differences in suggested interactions with rRNA are shown above G and H.

Both SURF6 and NPM1 have been proposed to interact with rRNA through weak promiscuous binding^12^ and we thus hypothesized that SURF6-NPM1 linkage occurs as a consequence of heterotypic interactions with rRNA, which are diluted upon NPM1 overexpression. To test whether heterotypic interactions with rRNA underly the thermodynamics of nucleolar assembly, we performed PITA following actinomycin D (ActD) treatment, which is known to halt transcription of nascent rRNA without affecting processing and assembly of pre-existing rRNA^19,20^ (**Fig. 3E**). As previously reported, the addition of ActD results in the progressive reduction of nucleolus size over the course of 4 hours (**Fig. 3F**, **S6**)^21^. Over time, the ΔΔG^tr^ of NPM1 and SURF6 increases, indicating weakened interactions relative to cells without ActD, consistent with NPM1 and SURF6 driving heterotypic phase separation through multivalent interactions with nascent, unfolded (or misfolded) rRNA transcripts, which become increasingly scarce under ActD treatment. Conversely, we find that the two r-proteins RPL23A and RPL5 display the opposite behavior, with their transfer free energies decreasing with the progression of ActD treatment (**Fig. 3G,H**), reflecting strengthened interactions that are consistent with specific binding to more fully processed rRNA.

These findings shed light on how heterotypic interactions driving phase separation facilitate sequential rRNA processing in ribosome biogenesis. Specifically, relatively nascent rRNA transcripts are available for more interactions with NPM1, SURF6, and other scaffolding components of the GC-matrix, compared to fully assembled ribosome subunits, providing a mechanism to facilitate the vectorial flux of processed subunits out of the nucleolus (**Fig. 4F**).

**Figure 4.**
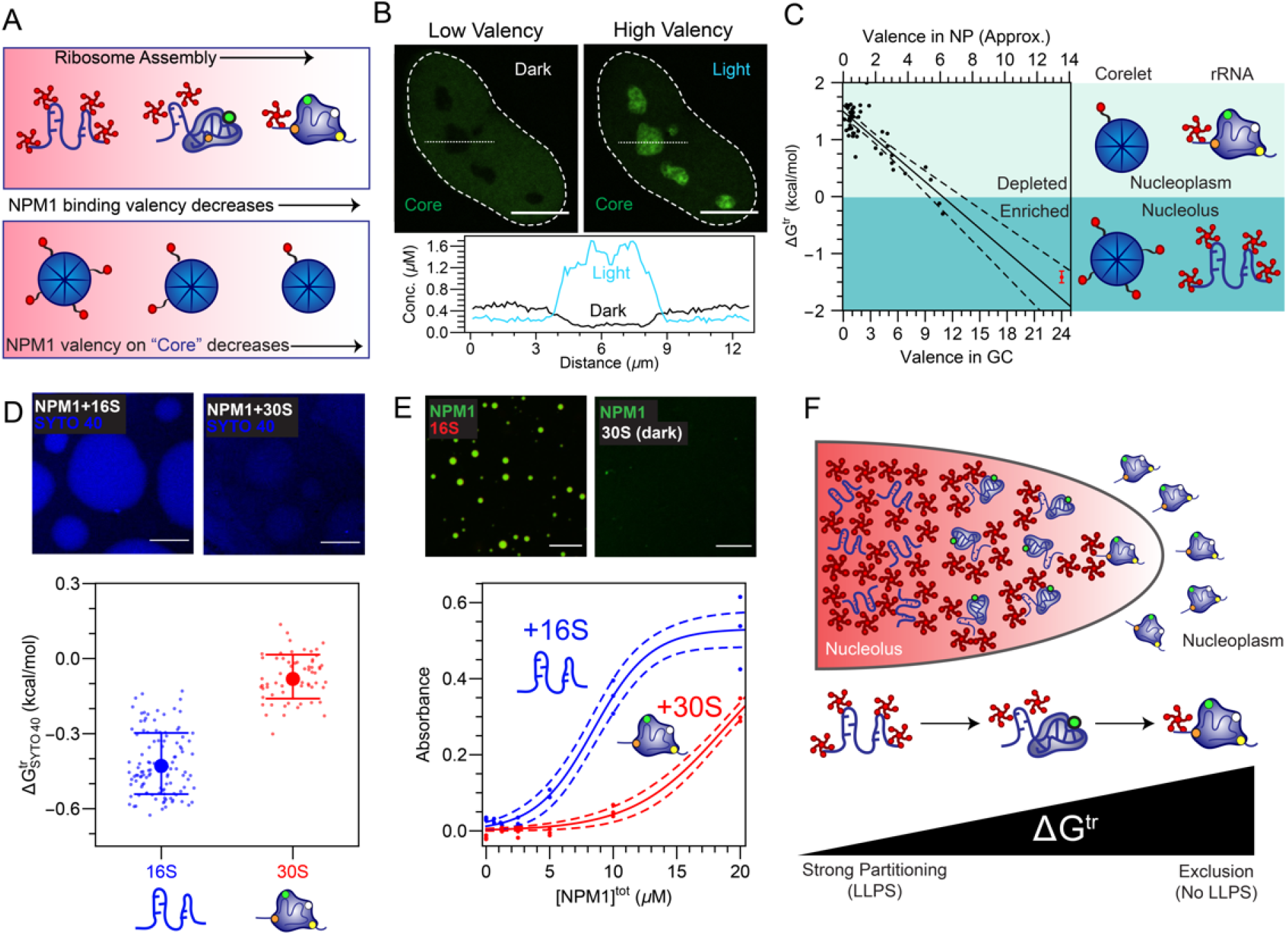
Composition-dependent heterotypic LLPS drives specific ribosomal subunit exclusion. (A) Top, schematic NPM1 valency as a function of rRNA folding/processing in the nucleolus and bottom, schematic of NPM1 valency on ferritin “cores” using the Corelet optogenetic system. (B) Images of a cell highlighting the partitioning of the “cores” (ferritin-iLID-GFP) before light (e.g. low effective valence) and after light (e.g. high effective valence) upon which NPM1-C binding sites on the core are saturated in this cell. Quantification is shown below corresponding to the dashed line shown in the images. (C) Corresponding quantification of the dependence for the ΔG^tr^ of the core as a function of the valence in the GC after light activation. Dotted lines are fits to data. (D) ΔG^tr^ measured by incubation with 6.5μM SYTO 40 to approximate the transfer free energies of 16S rRNA or the 30S small ribosomal subunit (at a total of 5μg/ml) into droplets formed with NPM1 (10μM) and 5% PEG droplets as indicated. Error bars represent standard deviation. Top pictures. (E) Turbidity assay of indicated concentration of NPM1 incubated with either the 16S rRNA or the 30S small ribosomal subunit in blue and green, respectively. 50μg/ml of the RNA species is added; validation of protein and RNA components in **Fig. S7**. 16S is labeled via a morpholino approach as described in the methods. Top, microscopy images with 10uM NPM1 with indicated RNA species. (F) Proposed mechanism for ribosomal subunit exclusion from the GC of the nucleolus driven by thermodynamics of nucleolar LLPS.

Indeed, binding of nascent transcripts by r-proteins eliminates multivalent binding sites for heterotypic scaffolding proteins, which could serve to effectively expel fully assembled preribosomal particles. We tested this concept using the biomimetic Corelet system, which consists of a 24-mer ferritin core, as a proxy for rRNA (**Fig. 4A**). Each ferritin subunit is fused to an optogenetic heterodimerization domain, which can be used to tune the effective valency of the particle with light^6^. We fused the cognate heterodimerization protein to an N-terminal truncated NPM1 (NPM1-C), which on its own partitions into nucleoli, with a ΔG^tr^ of approximately - 0.4kcal/mol (**Fig. S7**). In the absence of bound NPM1-C, the ferritin core is strongly excluded from nucleoli, with a ΔG^tr^ of approximately +1.4 kcal/mol (**Fig. S7**), consistent with large non-interacting assemblies being excluded from the nucleolus and other condensates^22–24^. However, upon increasing the valence of the core with light activation, its partitioning into the nucleolus increases, implying a more negative transfer free energy. This effect depends strongly on the valence of the Core: for valence <10 Corelets are excluded (ΔG^tr^>0), while for valence >10 Corelets are enriched (ΔG^tr^<0) within the nucleolus (**Fig. 4B,C, S7, and Supplemental Text**). This physical picture is supported by in vitro experiments with NPM1 droplets, which reveal that ΔG^tr^ is more strongly negative for 16S rRNA, compared to the 30S ribosomal subunit (comprised of 16S plus associated r-proteins **Fig. S7**) (**Fig. 4E**). Consistent with these measurements, in vitro phase separation of NPM1 is significantly weaker in the presence of the 30S subunit compared with 16S rRNA (**Fig. 4D**), underscoring how non-ribosomal protein bound (i.e. smaller) and highly solvent-exposed rRNAs are associated with favorable heterotypic interactions that promote partitioning and phase separation with nucleolar scaffold proteins (**Fig. 4D-E**). All together, these data suggest a mechanism that links the ability of nascent rRNA to promote phase separation and concentrate within nucleoli, while explaining how interactions with fully assembled ribosomal subunits are disfavored, leading to their thermodynamic exit from nucleoli.

Our findings lay the groundwork for establishing a quantitative understanding of the link between composition-dependent thermodynamics of condensate assembly and the free energy landscape of biomolecular complex assembly. In particular, we show that heterotypic biomolecular interactions give rise to high-dimensional phase behavior that yields C_sat_ values that vary with component concentrations, providing a mechanism for tuning condensate composition. This stoichiometric tuning enables “on demand” condensate assembly, such that phase separation only occurs in the presence of substrate, while simultaneously enabling a non-equilibrium steady-state flux of products (substrates), which are driven out of (or in to), the condensate during processing. This is likely relevant not only to the nucleolus, but also to many other phase-separated condensates that facilitate the formation of diverse biomolecular complexes, such as the spliceosome. Future work will exploit these intracellular thermodynamic design principles towards novel organelle engineering applications.

## Acknowledgements

We thank members of the Brangwynne laboratory for helpful discussions and comments on this manuscript. This work was supported by the Howard Hughes Medical Institute, the St. Jude Collaborative on Membraneless Organelles, and grants from the NIH 4D Nucleome Program (U01 DA040601), and the Princeton Center for Complex Materials, an NSF supported MRSEC (DMR 1420541). L.Z. was supported by the NSF graduate fellowship (DGE◻656466). R.W.K acknowledges support from NIH [R01 GM115634, R35 GM131891 and P30 CA021765 (to St. Jude Children’s Research Hospital)] and ALSAC. M.T. acknowledges support from NIH (F32 GM131524). Some images were acquired at the St. Jude Cell & Tissue Imaging Center, which is supported by SJCRH and NCI P30 CA021765; we would like to thank Drs. Victoria Frohlich and Jennifer Peters for their technical assistance. Data is available upon request.

## Author Contributions

J.A.R, L.Z., D.M.M., R.W.K. and C.P.B. designed research; J.A.R., L.Z., D.W.S., M.C.F., M.T. and M.W. performed research and contributed new reagents/analytic tools; J.A.R., L.Z., M.C.F., M.T., and D.M.M. analyzed data; J.A.R, L.Z., and C.P.B. wrote, and all authors reviewed and edited the paper.

## Methods

### Preparation of mammalian cells

NPM1 (Sino Biological), SURF6 (DNASU), RPL5 (IDT Gene Block synthesized), RPL23A (DNASU), DCP1A, and Coilin were cloned into the FM5-mCherry, FM5-GFP, or FM5-EYFP plasmids. DNA fragments encoding the proteins of interest were inserted into the restriction digested linearized FM5 backbones using In-Fusion Cloning Kit (Takara). The resulting constructs were fully sequenced to confirm the absence of unwanted substitutions.

Cell lines were created through lentiviral transfections of ATCC HeLa cells using Lenti-X cells transfected with Fugene HD transfection reagent. HeLa cells are plated on fibronectin-coated, glass dishes prior to imaging. Cell lines were maintained at standard conditions (37°C, 5% CO2).

### Imaging of mammalian cells

Cell lines were imaged on a Nikon A1 laser scanning confocal microscope using an oil immersion objective, Plan Apo 60×/1.4. Imaging conditions (i.e. gain and laser intensity) were optimized to increase signal to noise within the dynamic range. Consistent intensity referencing including, when applicable, conversion into a concentration were done as previously reported^6^. When two proteins are imaged concurrently, sequential scanning was used. OptoG3BP1 experiments were done as previously reported.^4^

### Actinomycin D treatment

Cells were treated to a total concentration of 1ug/mL actinomycin D (ActD, sigma) made from a 0.5mg/mL stock solution) in DMEM media and maintained at standard conditions before and throughout imaging.

### Arsenite treatment on G3BP1 or optoG3BP1 expressing cells

G3BP1/2 Knockout Cells^15^ expressing, via lentiviral transfection and integration, protein constructs (either G3BP1 or optoG3BP1) were treated to a total concentration of 400uM sodium arsenite for 1 hour prior to imaging and maintained at standard conditions before and throughout imaging. For optoG3BP1 experiments, we cloned Cry2 N-terminal of G3BP1ΔNTF2 similarly as done in^25^.

### Methods for the determination of C^dil^, C^den^, C^tot^, ΔG^tr^, and fits from *in vivo* images

To determine the ΔG^tr^ from a confocal image, one must determine the dilute and dense phase concentrations. For nearly all data, a representative area of the dense phase (e.g. GC of nucleoli, stress granules, etc.) was determined as the average of the computed most intense box of size 7 by 7 pixel units (in rare cases, 5 by 5 pixels was used when droplets were small), a representative area of the dilute phase was defined by a box of size 21 by 21 pixels nearby this intense box was manually chosen (to minimize any bias by out of plane droplets), and the background intensity was found by computing the least intense box of 101 by 101 pixels. The concentrations were calculated by subtracting the average intensity in the respective phase from the background intensity and applying the conversion into a concentration as previously reported^6^. In some cases (e.g. Coilin) two separate images were taken to ensure that both the dilute and dense phase were both within the dynamic range. For each construct, a manual minimum dense phase concentration was chosen to eliminate cells without droplets or with too small droplets. When calculating C^tot^ for NPM1, a custom Mathematica script was applied that segmented the nucleus, using the intensities as the background and nucleoplasmic as inputs, and reported the average intensity being an approximate for C^tot^. When calculating C^tot^ for G3BP1, a 41 by 41 box around the location of a stress granule, as described above, was used as an approximate for C^tot^. For optoG3BP1, the C^tot^ was given by the pre-activation intensity in the region described by the dilute phase. Fits throughout the text are fit in Mathematica using the “NonlinearModelFit” function to the empirical form 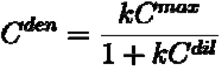 and lines shown throughout the text are the mean prediction bands given by the Mathematica function “MeanPredictionBands”. For fitting relationships involving the total concentration for NPM1, we assume that the dependence of the area fraction is linear within our experimental changes, this dependence is used to for the dependence of C^den^ and C^dil^ on C^tot^ (and vice versa). To plot ΔG^tr^ vs. C^dil^, the data were transformed by 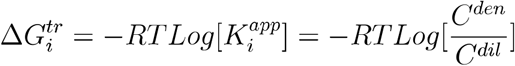 where T = 37°C or 310.15K (or room temperature for the *in vitro* data). The exception to this fit is for the Corelet data where the trend of ΔG^tr^ vs. valence which is simply fit with a line in Mathematica with “MeanPredictionBands” similarly shown.

### Methods for the quantification of and theoretical considerations for ΔΔG^tr^

To quantify ΔΔG^tr^ for component i as a function of component j (i.e. 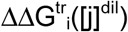) we expressed both components i and j and calculate concentration and partition coefficients as described above. In these experiments, we attempt to either minimize the amount of component i and/or keep [i]^dil^ < [j]^dil^. To correct for changes in the concentration of [i]^dil^ as the volume fraction of the dense phase and/or at different over expressions of i, we establish a reference curve for 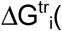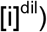 without increasing the concentration of j and fit this curve as described in the previous section. Using this reference curve, we preform the calculation of ΔΔG being 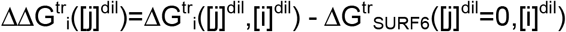 We typically fit this trend to a linear or quadratic fit and display the corresponding mean confidence bands in Mathematica as described previously. The assumption we are utilizing for this referencing assumes that the effect of i and j addition on the transfer free energy is additive within this range.

### Purification of non-ribosomal proteins

Recombinant poly-histidine tagged full-length wild-type and C21T/C275T mutant NPM1 were expressed in BL21 (DE3) *E. coli* cells (Millipore Sigma, Burlington, MA) and purified as previously described^17^. Briefly, cells were lysed by sonication and proteins were isolated from the insoluble fraction using Ni-NTA affinity chromatography. Affinity tags were removed via proteolytic cleavage with TEV and further purified on a C4 HPLC column (Higgins Analytical, Mountain View, CA) and lyophilized. Lyophilized NPM1 C21T/C275T was resuspended in 20 mM Tris containing 6 M GuHCl pH 7.5 and labeled overnight with Alexa-488 maleimide dye (Invitrogen) at 1:1.5 (protein:dye). Excess fluorescent dye was removed by dialysis. Unlabeled NPM1 WT and fluorescently-labeled NPM1 C21T/C275T (NPM1-488) in 20 mM Tris, 6M GuHCl, 2 mM DTT pH 7.5 were mixed at 9:1 ratio and were refolded by dialyzing overnight in 10 mM Tris, 150 mM NaCl, 2 mM DTT, pH 7.5 buffer. Aliquots of NPM1 were flash frozen and stored at −80 °C.

Poly His-tagged SURF6-N with a TEV cleavage site were expressed in BL21 Rosetta 2 (DE3) *E. coli* cells (Millipore Sigma, Burlington, MA) and isolated from inclusion bodies. Cells were lysed by resuspension in 50 mM sodium phosphate 300 mM NaCl 0.1% Triton X-100 pH 8.0 buffer. The pellet fraction was collected and purified on a Ni-NTA column under denaturing conditions. His-tagged SURF6-N was treated with TEV protease to cleave the tag and further purified using a C4 HPLC column. Lyophilized SURF6-N was resuspended in 20 mM Tris, 6 M guanidine HCl, 2mM DTT pH 7.5 and dialyzed in high salt buffer (10 mM Tris, 1 M NaCl, 2 mM DTT, pH 7.5) and stored in small aliquots at −80 °C. These SURF6-N stocks were diluted with 10 mM Tris, 2 mM DTT, pH 7.5 to give final salt concentration for assays of 150 mM NaCl.

### Purification of ribosomal components

All components were purified from *E. coli* K12, strain A19 grown in Lauria Broth. Cultures were grown at 37 °C in an InnOva 4230 incubator/shaker (New Brunswick Scientific) to an OD_600_ of ~0.8 and harvested by centrifugation. Cell pellets were suspended in 20 mM Tris-HCl, 50 mM MgOAc, 100 mM NH_4_Cl, 1 mM TCEP, 0.5 mM EDTA, pH 7.5 and lysed using high pressure homogenization. The soluble fraction was layered onto a 30% sucrose cushion prepared in buffer A [10 mM Tris-HCl, 6 mM MgOAc, 50 mM NH_4_Cl, 1 mM TCEP, 0.5 mM EDTA, pH 7.5] and centrifuged at 30,000 rpm for 16 hours at 4 °C in a SW 32 Ti Swinging-Bucket Rotor (Beckman Coulter). The resulting pellet was gently suspended in buffer A and subjected to two additional rounds of purification on a 30% sucrose cushion. After these steps, the sample was purified using a 10-35% linear sucrose gradient prepared in buffer A and centrifuged at 21,000 rpm for 20 hours at 4 °C. Samples containing pure 70S ribosomes were pooled and centrifuged at 30,000 rpm for 16 hours at 4 °C. The resulting pellet was suspended in buffer A for separation into its constituent ribosomal components.

Ribosomes in buffer A were dialyzed overnight against 4 liters of buffer B [10 mM Tris-HCl, 1 mM MgOAc, 100 mM NH_4_Cl, 1 mM TCEP, 0.5 mM EDTA, pH 7.5] to allow dissociation into 50S and 30S ribosomal subunits. The resulting solution was applied to a 10-35% linear sucrose gradient prepared in buffer B and centrifuged at 21,000 rpm for 18 hours at 4 °C. Fractions containing 30S small ribosomal subunit were pooled and centrifuged at 30,000 rpm for 16 hours at 4 °C. The pellet was gently suspended in 10 mM Tris, 150 mM NaCl, 2 mM DTT, pH 7.5 for use in turbidity and microscopy measurements.

16S rRNA was extracted from the 30S small ribosomal subunit using standard phenol:chloform extraction. 16S rRNA was then purified to homogeneity by 1.2% native agarose electrophoresis, gel excision, and electroelution. The sample was washed into 10 mM Tris, 150 mM NaCl, 2 mM DTT, pH 7.5, using a Millipore Amicon Ultra-15 centrifugal filter device. RNA purity was confirmed using a 1.2% denaturing 1× TBE denaturing agarose gel.

### Determination of partition coefficients for NPM1 and SURF6-N in *in vitro* droplets with wheat germ rRNA

Multicomponent *in vitro* droplets were prepared on chambered coverglass slides coated with Repel Silane ES and PF-127. Droplet mixtures contained 5 μM of NPM1 and SURF6-N (with 500 nM SURF6-N-A647) and 25 ng/μL wheatgerm rRNA (bioWORLD, Dublin, Ohio, U.S.A.) in 10 mM Tris, 150 mM NaCl, 2 mM DTT buffer pH 7.5. Wheatgerm RNA was purified using phenol:chloroform extraction. Titrations were performed by adding equal volumes of NPM1 (with 240 nM NPM1-Alexa 488) over a range of concentrations (0.5 μM to 18.4 μM) to the droplet solutions. Single-component droplets were prepared by mixing NPM1-Alexa 488 or NPM1-Alexa 594 with buffer containing 5% PEG (MW=8000 Da). Fluorescence microscopy imaging was performed using a Zeiss LSM 780 NLO point scanning confocal microscope with 63X oil immersion objective (NA 1.4). Droplet mean intensities were analyzed using FIJI software using the default threshold settings. From the mean fluorescence intensities, the apparent partition coefficients (Kapp) of SURF6-N-Alexa 647 and NPM1-Alexa 488 were determined. Kapp for SURF6-N-Alexa 647 and NPM1-Alexa 488 were calculated from the equation: Kapp = (*I*_DP_−*I*_bkrd_)/CF]/(*I*_LP_−*I*_bkrd_), where *I*_DP_ is the average of mean intensities of droplets, *I*_LP_ is the mean intensity of the regions outside the droplets, *I*_bkrd_ is the mean intensity of an image of buffer alone and CF is the quantum yield correction coefficient for the fluorescent dye (Mitrea, *et al.*, 2018 *Nature Communications*). To determine the effect of increasing SURF6-N concentrations on multicomponent droplets a similar titration was performed over a range of SURF6-N concentrations containing 240 nM SURF6-N-A647.

### Turbidity assays with 16S rRNA and 30S small ribosomal subunit

Solution turbidity was used to detect phase separation of NPM1 with 16S rRNA and the 30S small ribosomal subunit. All assays were performed and measured in triplicate in a reaction volume of 10 μL in 10 mM Tris, 150 mM NaCl, 2 mM DTT, pH 7.5. For all reported assays, turbidity measurements were performed at room temperature and measured at 340 nm on a NanoDrop 2000C (Thermo Scientific, Waltham, MA, USA) 10 minutes after sample mixing. For each condition examined, the ribosomal component (normalized to 50 μg/mL RNA) was mixed with serially diluted NPM1. The corresponding images were acquired using a Zeiss LSM 780 NLO point scanning confocal microscope with 63X oil immersion objective (NA 1.4). Droplet mixtures contained 10 μM of NPM1-Alexa 488 and 50 μg/mL 16S rRNA or 30S small ribosomal subunit normalized to 50 μg/mL RNA. All imaging was performed 30 minutes after mixing at room temperature in 10 mM Tris, 150 mM NaCl, 2 mM DTT, pH 7.5 in chambered coverglass slides coated with Repel Silane ES and PF-127. To detect 16S rRNA localization, the 16S rRNA was mixed with an equimolar amount of a 13mer morpholino complementary to nts 110-122 (Gene Tools, LLC) labeled with Alexa 555 (Life Technologies) prior to NPM1 addition. The morpholino used was labeled via its 5’ amine using with an Alexa Flour succinimdyl ester dye conjugate according to the manufacturer’s protocol. Excess dye was removed thru dialysis. NPM1 was labeled as described above.

### Determination of partition coefficients for 16S rRNA and 30S small ribosomal subunit in *in vitro* droplets

Pre-equilibrated, single-component droplets of 10 μM NPM1 (with 100 nm of NPM1-Alexa-594 and 5% PEG in 10 mM Tris, 150 mM NaCl, 2 mM DTT, pH 7.5) were used to measure differential partitioning of 16S rRNA and the 30S small ribosomal subunit. Droplets were allowed to equilibrate prior to addition of 16S rRNA or 30S small ribosomal subunit; both species were normalized to 5 ng/uL RNA concentration. SYTO 40 (Invitrogen) was used to detect 16S rRNA or RNA within the 30S small ribosomal subunit. Imaris software (Bitplane) was used to measure mean fluorescence intensities for these droplets. Kapp for 16S rRNA and 30S small ribosomal subunit were calculated from the equation: Kapp = (*I*_DP_−*I*_bkrd_)/(*I*_LP_−*I*_bkrd_), where *I*_DP_ is the average of mean intensities of droplets, *I*_LP_ is the mean intensity of the regions outside the droplets, *I*_bkrd_ is the mean intensity of an image of buffer alone.

### Application of Flory-Huggins Theory

Flory-Huggins theory was used to approximate the behavior of multicomponent phase separating polymers^13^. Under this theory, the chemical potentials for each component are:

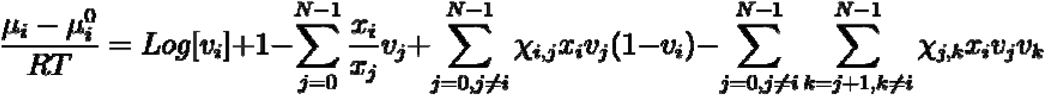

Where *μ*_*i*_, *x*_*i*_, and *v*_*i*_ are the chemical potential, the number of segments (e.g. amino acids) per molecule, and the volume fraction for component *i* respectively. *X*_*ij*_ is the interaction intensity per segment (defined here ‘per segment’ which is slightly different than Flory did) on the polymer chain determined from a lattice model of *z* dimensions as 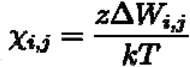 where *W*_*i,j*_ is the energy of each contact between components *i* and *j* and 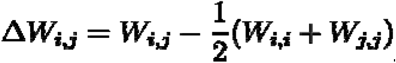 To demonstrate this and other principles, we make three component phase diagrams (**Fig. S3**). For simplicity RT=1. The solvent (i.e. component ‘0’) has x=0 whereas, for simplicity, the two polymer components are set as x=1000. The polymers are set as χ=0.54811 yielding a partition coefficient for each in water (e.g. homotypic) as 50. For the strength of the heterotypic interactions relative to homotypic ones, χ is set to 0.0, −0.04, 0.015 for heterotypic being equal, greater, or less then homotypic interactions, respectively. Analytical solutions are found via Mathematica using the FindMinimum function minimizing the difference in the chemical potential in the dilute and dense phases for all three components and progressing along the phase diagram by incrementally moving along nearby solutions and constraining the change in the variables to between 25% and 175%.

---

**Figure S1.**
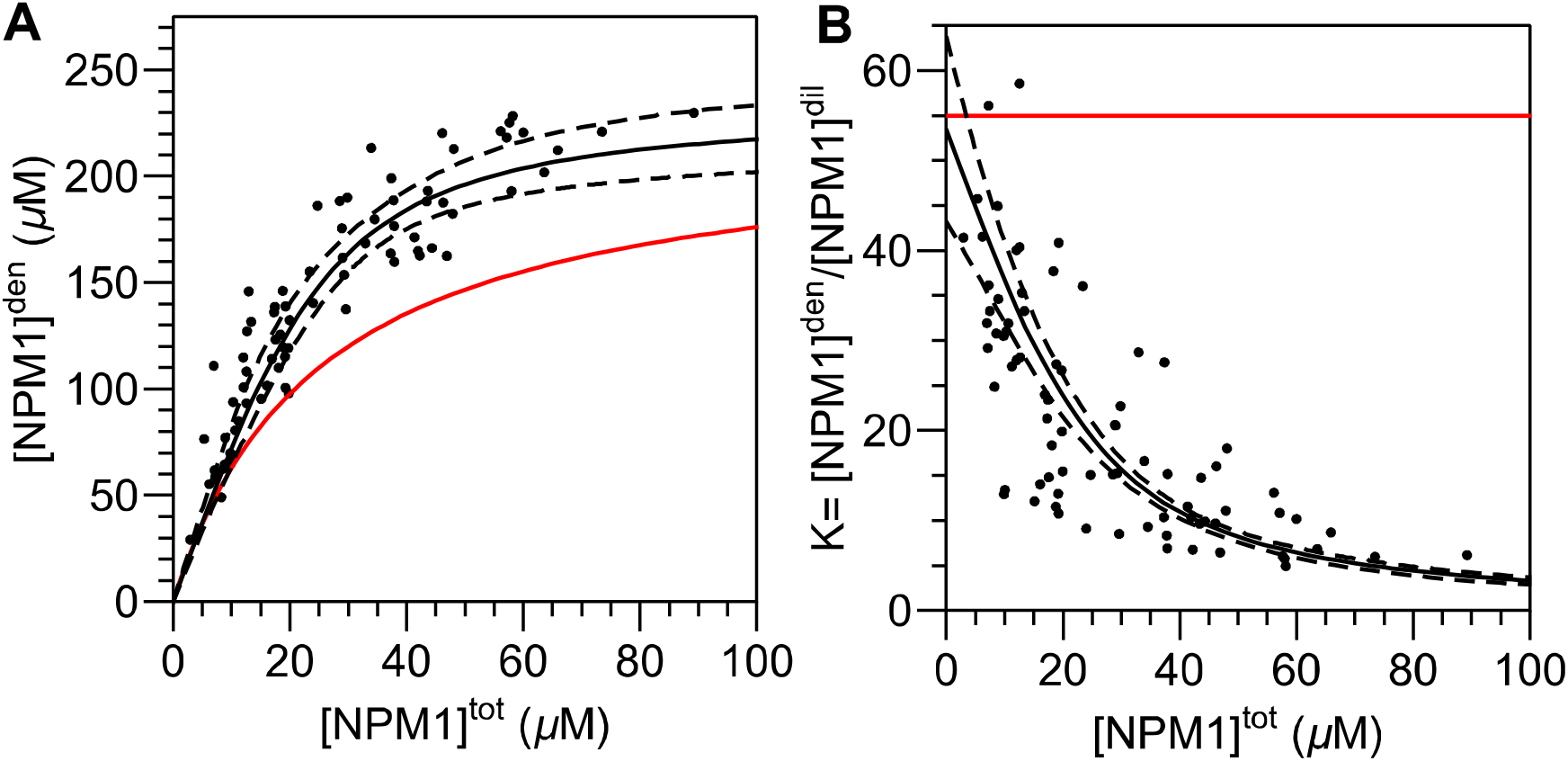
NPM1 lacks a fixed C^dil^ and C^den^ suggesting Nucleoli undergo multicomponent mediated phase separation. (A-B) Dependence of NPM1 total overexpression in the nucleus vs. (A) its measured concentration in the relevant dense phase (‘den’ being the GC of nucleoli) or (B) its apparent partition coefficient being the ratio of its concentration in the dense to its concentration in the dilute phase. Dashed Lines represent mean confidence intervals to fits described in the methods. Red lines represent expected trends for single component phase separation.

**Figure S2.**
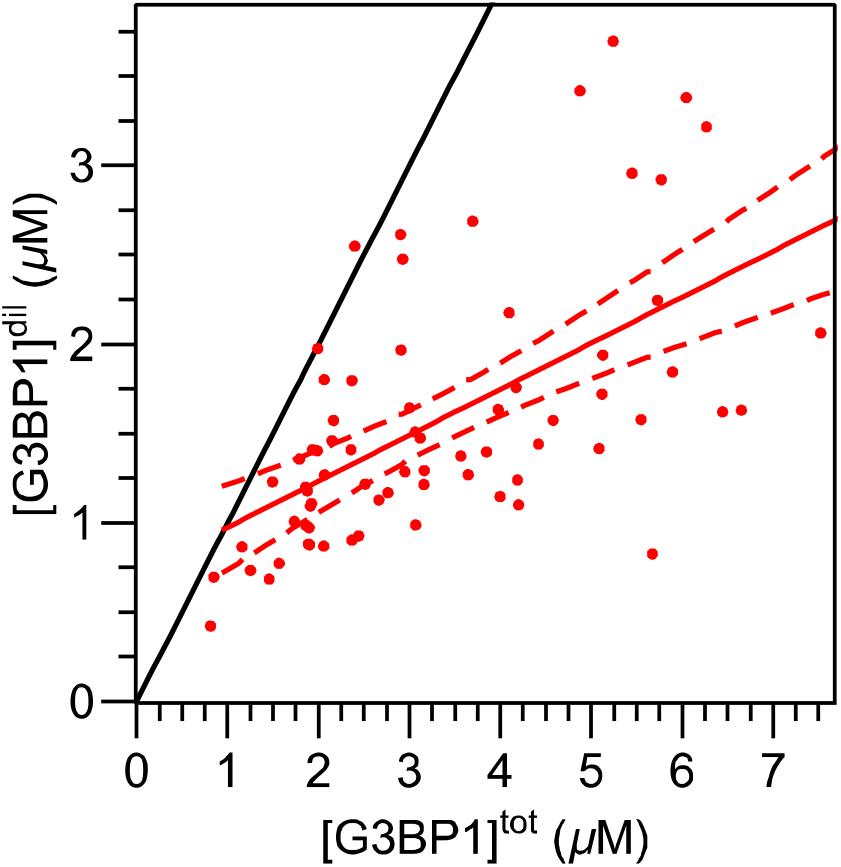
Similar to optoG3BP1, no fixed C_sat_ exists for G3BP1. Dependence of the concentration of approximated total cytoplasmic concentration of G3BP1 as a function of the dilute (i.e. cytoplasmic) concentration in G3BP1/2 knockout cells after arsenite treatment. Dashed Lines represent mean confidence intervals to fits described in the methods.

**Figure S3.**
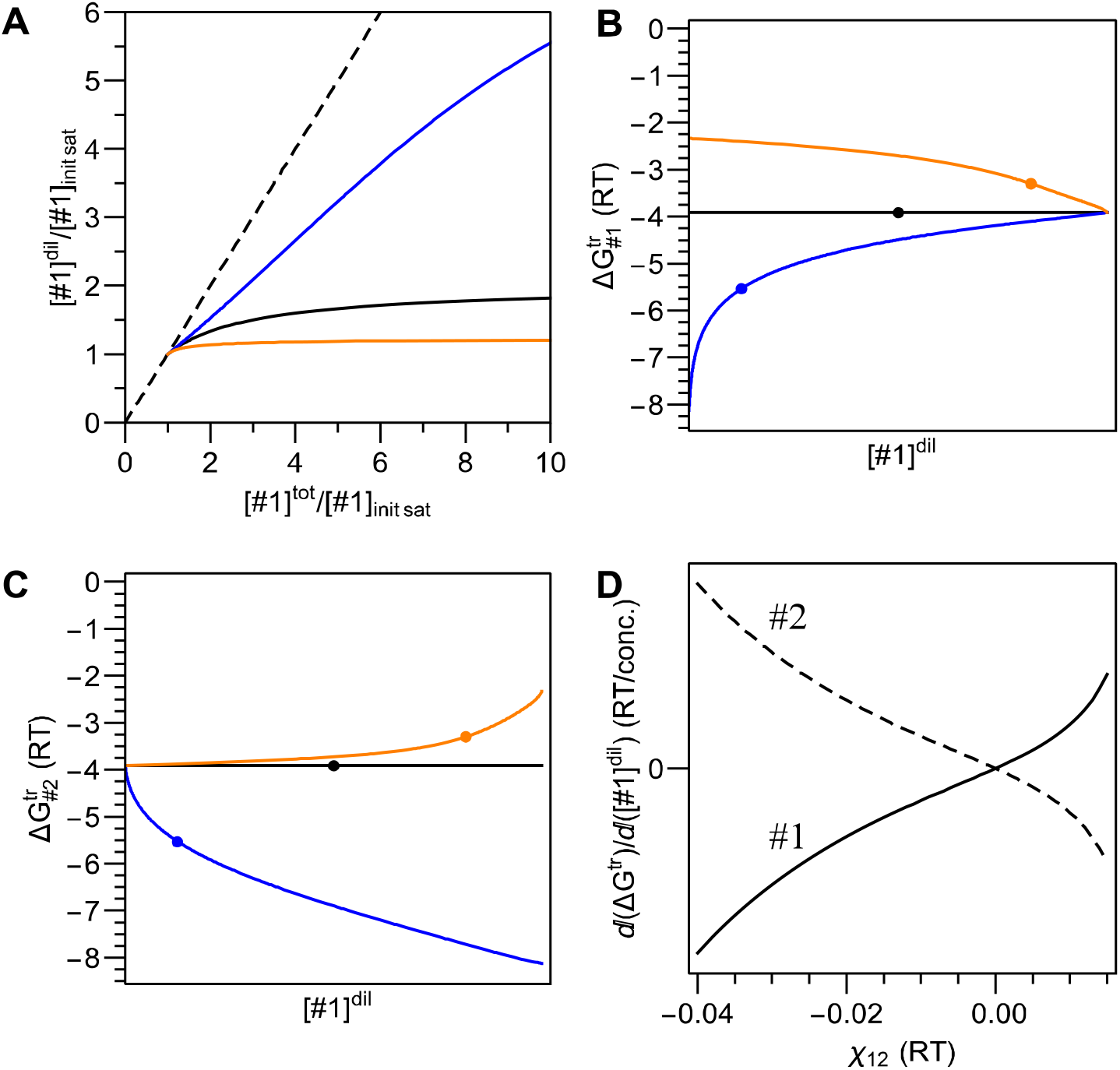
*In silico* validation of PITA using Flory Huggins theory. Phase separation of two (non-solvent) components, denoted #1 and #2, with their heterotypic interactions being equal, stronger, and weaker, then their homotypic interactions shown as black, blue, and orange, respectively for A-C. Note the dilute phase in the bottom left corner of the plot. (A) The initial dependence of the [#1]^dil^ on [#1]^tot^ at fixed [#2]^tot^ such that phase separation will occur at the ‘goldilocks point’ being when [#1]^tot^=[#2]^tot^. Note the axis are normalized by the initial saturation (init sat) concentration, i.e. lowest [#1]^tot^ where phase separation emerges. The dashed line is the 1:1 line where expected without phase separation. (B,C) Dependence on ΔG^tr^_#1_ (D) or ΔG^tr^_#2_ (E) as a function of [#1]^dil^. Circles indicate the location of the ‘goldilocks point’. (D) Dependence of the change in ΔG^tr^ with respect to [#1]^dil^ as a function of the heterotypic interaction strength χ_12_ (where more negative implies stronger heterotypic interactions) at the ‘goldilocks point’ for the transfer free energy of #1 and #2 as indicated.

**Figure S4.**
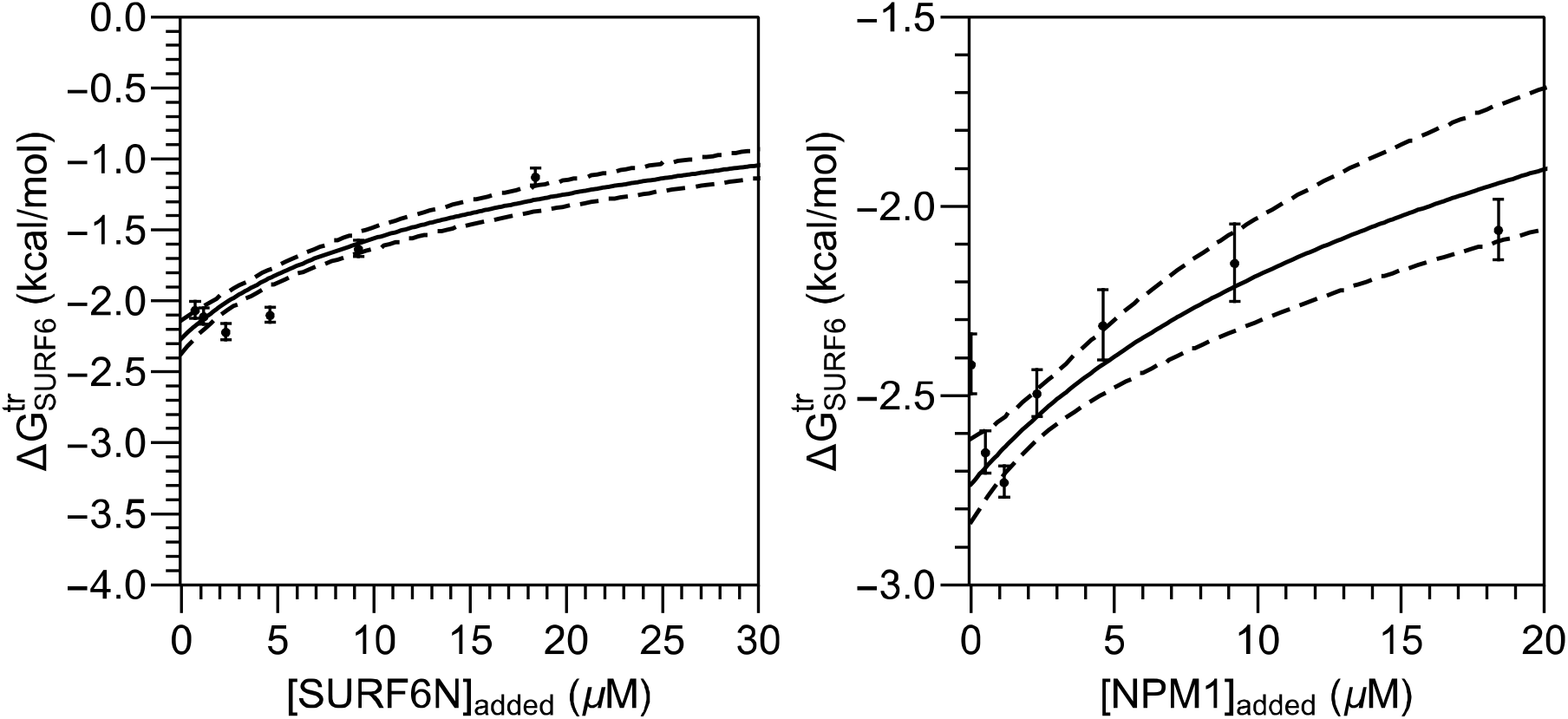
*In vitro* destabilization of SURF6N partitioning by overexpression of itself or NPM1. Changes in the transfer free energy of SURF6N into multicomponent droplets as additional SURF6N (A) or NPM1 (B) is added on top of NPM1:SURF6N:rRNA ternary droplets as described in the methods.

**Figure S5.**
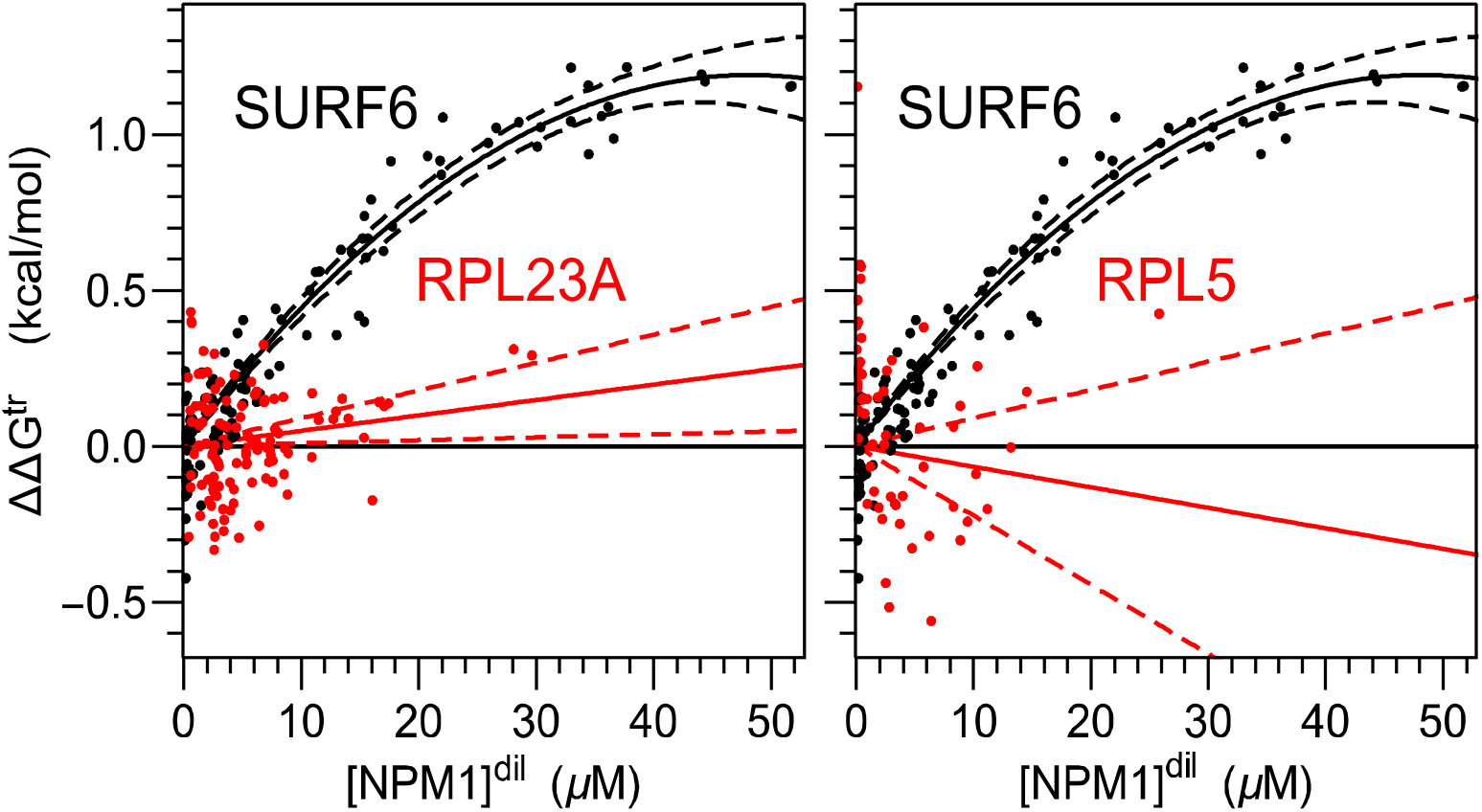
R proteins and NPM1 ΔΔG^tr^. Change in the transfer free energy of r-proteins RPL23A and RPL5 compared to the SURF6 and with as NPM1concentration is increased.

**Figure S6.**
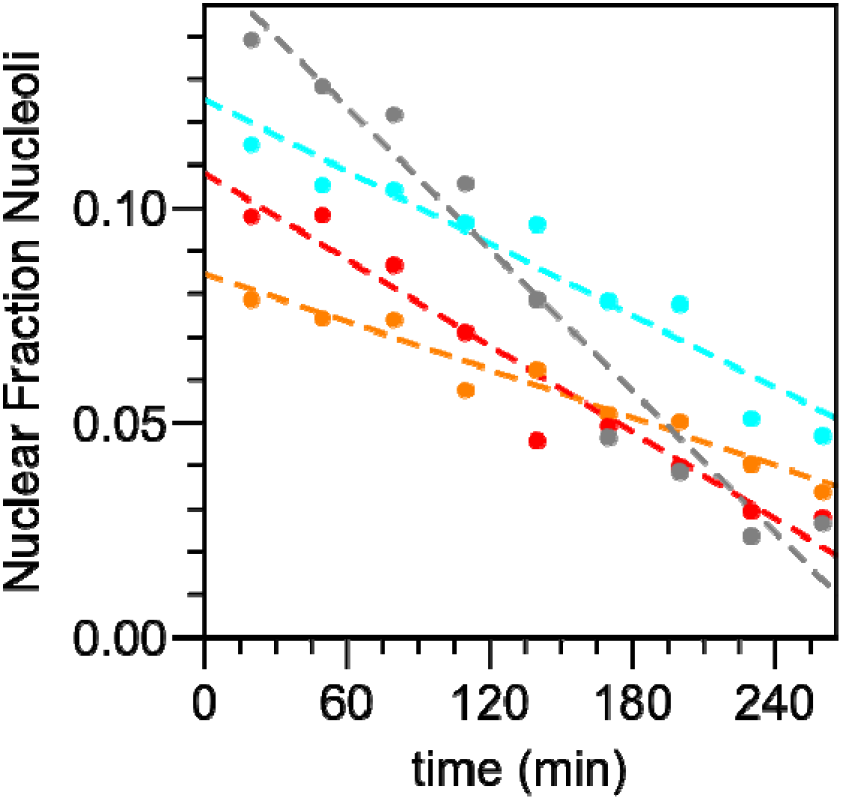
ActD decreases nucleolus size. Nucleolar fraction of image area as a function of time after addition of ActD in individual cells expressing NPM1-mCherry. Colors indicate same cells as in Fig. 3G, right.

**Figure S7.**
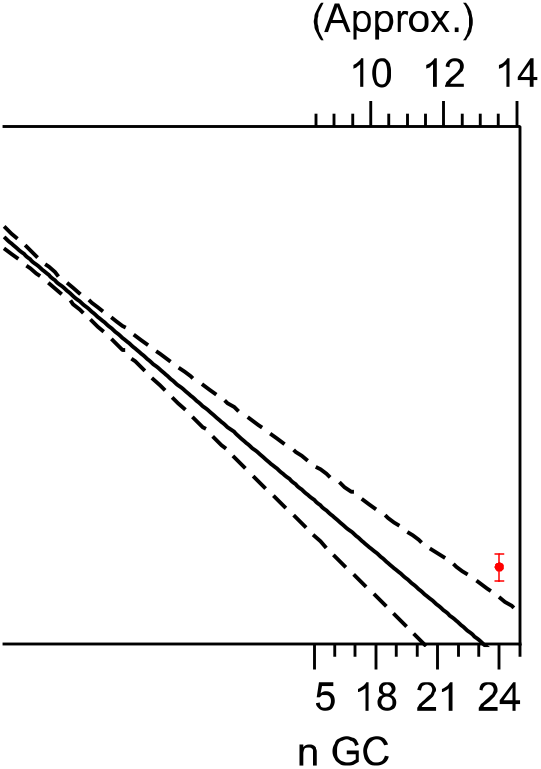
Characterization of Corelet non-ideality and extrapolation from high valence. (A) Dependence on the transfer free energy for the N-terminal half of NPM1 (NC)-sspB in cells without the core expressed (orange being a ΔG^tr^) or with the indicated valences following core activation (black being a 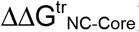). The 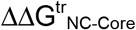 in this case is the energetic difference between the NC and core channels which is approximately the energetic difference for transferring an additional NC to the core at that valence. (B) At valences higher then 24, the transfer free energy is approximated as quadratic and extrapolated back to a valence of 24 to obtain the transfer free energy at this valence. (C) Transfer free energy reported from the sspB channel as a function of valence which is weighted by the number of sspB molecules (due to the number of mCherry molecules observed being proportional to each molecule’s valence as opposed to the core where it is always constant at 24 GFPs).

**Figure S8.**
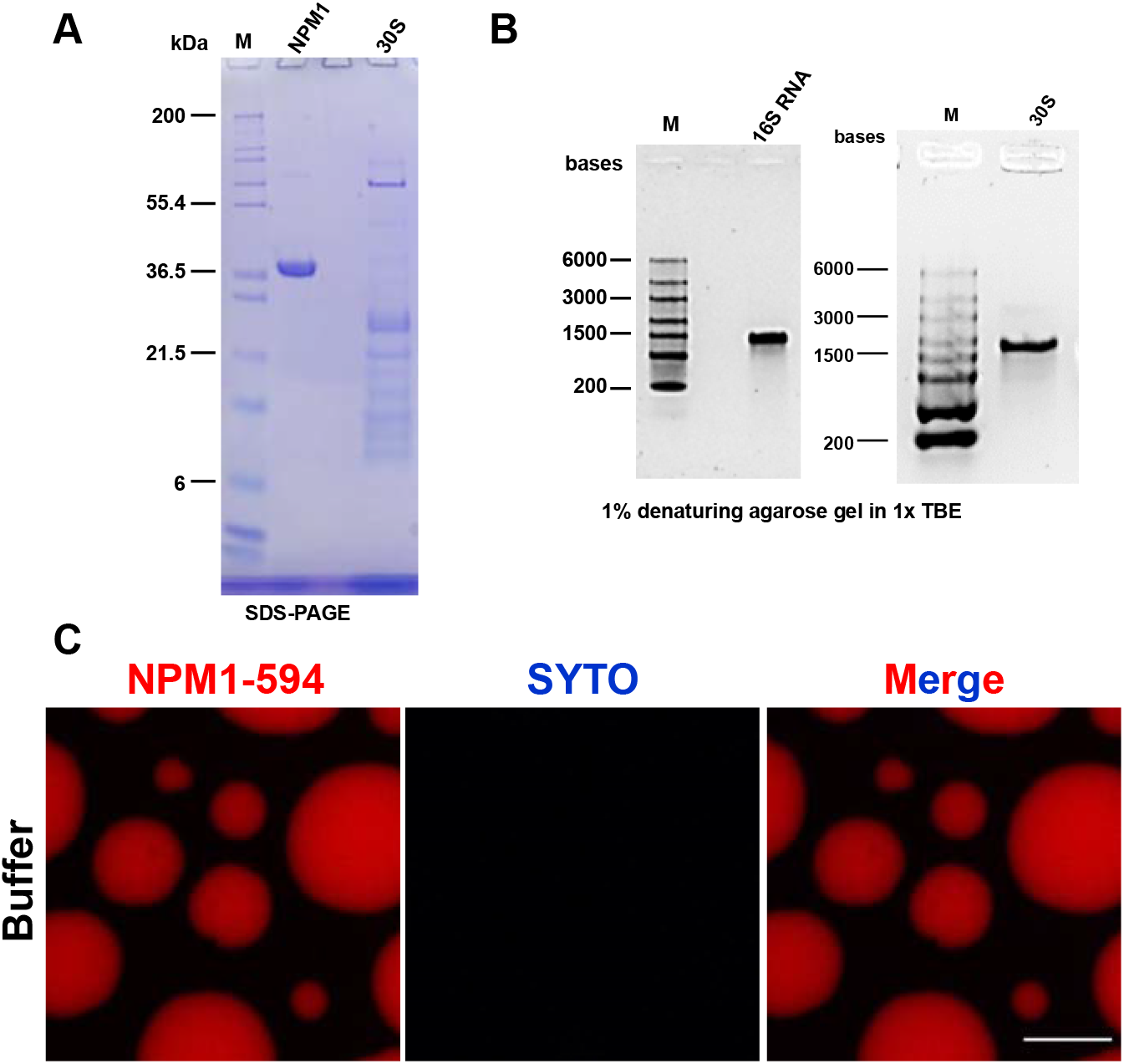
Controls for ribosomal mimics. Representative SDS-PAGE (A) and denaturing agarose gel (B) confirming purity of reagents used in experiments presented in Figure 4d and e. (C) Representative microscopy images for 10mM NPM1-594 droplets formed with 5% PEG without any rRNA showing limited florescence indicating neither NPM1 nor PEG binds SYTO 40 and the droplet environment does not promote florescence of SYTO 40.

## Supplemental Text

### Clarification on component number for phase separation terminology

Throughout this paper we make a few notation choices for the purpose of clarity to a non-specialist reader and for a reasonable balance of brevity and logical parallel statements. In particular, our replacement of the classical term “two-component phase separation”, where one component is the solute (typically, a polymer) and the other is the solvent (typically water), with the term “single component phase separation” where the single component is referring to the single biopolymer undergoing phase separation from the mean field bulk solvent of the cell. This choice is made to contrast a single biopolymer from a case where multiple biopolymers undergo phase separation which we refer to as multicomponent phase separation; thus, these terms set up the logical dichotomy between single component and multicomponent phase separation widely discussed throughout our paper. This notation is consistent with the conceptual perspective where polymer phase separation is analogous type of gas to liquid condensation where the solvent is ignored.

### Quantification of Stability Between Two Phases

Central to quantifying the thermodynamics of interactions which determine the relative abundance (i.e. partitioning) between two phases is the identification of a parameter derived from experimental observables and determination of its thermodynamic meaning. To derive such a thermodynamic parameter, we begin with the definition that two phases (dilute and dense abbreviated as “dil” and “den”, respectively) at equilibrium will have equal chemical potentials in each phase, for every component i:

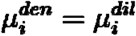

Specifying the component activity, *a*, in each phase, this expression can be written as:

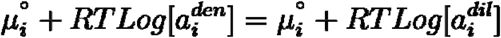

Here, the standard state or ideal state chemical potential 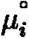 is equivalent in both phases (cite flory textbook), with the reference ideal solution being pure water for both phases. The activity is typically defined as 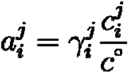 with 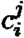 being the molar concentration of component *i* in phase *j*, *c*° being the standard reference concentration of 1 molar although all properties are extrapolated from infinite dilution, and 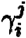 is the activity coefficient to account for non-idealities from a solution where non-solvent components are negligible (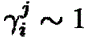 see further discussion at the end of this section for a more in depth discussion on 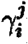). Applying these definitions and simplifying further:

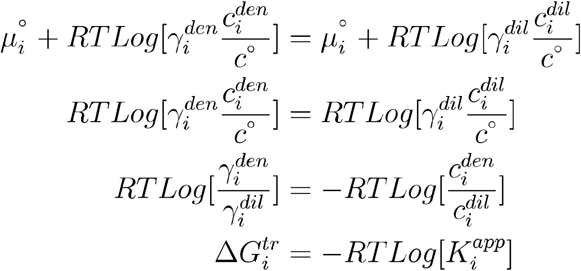

Thus, in this case, the ratio of the concentration between the dense and dilute phases of component i, the apparent partition coefficient, 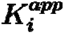 is equal to the energetic difference in non-ideality of the solution between the two phases. More generally, this energy is the transfer free energy as it is the free energy difference for a standard state concentration of molecules being transferred from the dilute to the dense phase.

We note, however, that our use of the activity coefficient includes both non-specific and site-specific interactions (described in^26^). Although beyond the scope of this paper, we note the existence of thermodynamic formalisms to separate these interactions^26–28^.

### Expected C_sat_ for NPM1

The total concentration of NPM1 in the whole cell has been measured as ~7μM^29^. By assuming that the total cell to nuclear volume is roughly 3.5 fold and because we measure its concentration in the cytoplasm to be negligible, we obtain a total nuclear concentration of NPM1 of ~25uM. Using the apparent partition coefficient (~55) and nucleolar fraction (~0.1) extrapolated to zero expressed fluorescent NPM1, one can use conservation of particles to approximate the nucleoplasmic concentration of endogenous NPM1 to be ~4uM. Thus, if this value were fixed at a C_sat_ then we would expect that the total (endogenous + florescent) NPM1 in the nucleoplasm to remain at this C_sat_ and thus the total concentration of expressed fluorescent NPM1 (c^tot^) would determine the dilute concentration of NPM1 (c^dil^) via the formula 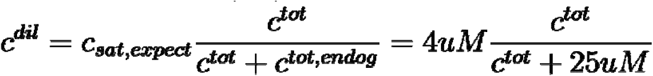 as shown. We note that while these values are only approximate, the general curvature in **Fig. 1B**is not consistent with this expected trend providing additional evidence that Nucleoli are not governed by a fixed C_sat_ for NPM1. We do not claim that this curvature will be general, as some systems may approach a C_sat_ for the over expressed component as the heterotypic interactions become diluted; approaching an effective C_sat_ with overexpression will likely be determined by the effective strength of homotypic interactions relative to the heterotypic ones.

### Basic logic for heterotypic stabilization

The theoretical basis for the correlation between the strength of heterotypic interactions in changing in the partition coefficient can be understood by expanding the difference of chemical potential of component i between the two states focusing on the difference between the interaction between component i and a non-solvent component m; proceeding:

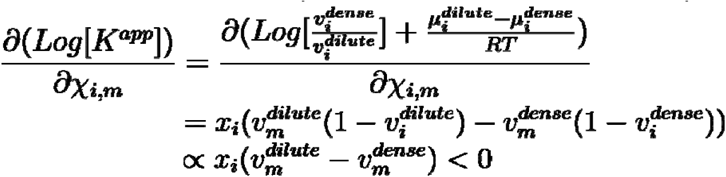

Thus when heterotypic interadctions are stronger than homotypic ones (i.e. *x*_*i,m*_ < 0), then the partition coefficient will tend to increase whereas when they are weaker, the partition coefficient will tend to decrease.

